# Validation of a CRISPR/cas9-based technology platform for examining specific immune gene functions in an experimental murine model of IBD

**DOI:** 10.1101/619791

**Authors:** Rui Wang, Sean Graham, Ning Sun, Donna McCarthy, Ruoqi Peng, Jamie Erickson, Xiaochun Zhu, Marc Wurbel, Robert Dunstan, Namjin Chung, Edda Fiebiger, Tariq Ghayur, Jijie Gu

## Abstract

Inflammatory bowel diseases (IBD) are complex, multifactorial disorders characterized by chronic relapsing intestinal inflammation. Association studies have identified hundreds of genes that are linked to IBD and potentially regulate its pathology. The further dissection of the genetic network underlining IBD pathogenesis and pathophysiology is hindered by the limited capacity to investigate the role of each GWAS association through functional studies, including the generation of knockout animal models for each of the associated genes. The CRISPR/Cas9 system represents a cutting edge technology which has the potential to transform the field of IBD research by facilitating the introduction of genetic alterations in an efficient and effective manner. Using the CD40-mediated-colitis model, our results demonstrate the validity of a CRISPR/Cas9-based platform as a tool for the validation of target genes or interference strategies in experimental IBD. The utilization of this discovery strategy will allow for the timely *in vivo* validation of therapeutic targets as the rapidly emerge from current genetic and genomics efforts with human disease tissue. As such, the CRISPR/Cas9-based platform can significantly shorten the time span between target identification and generation of proof of principle experiments for drug discovery.

## Introduction

Inflammatory bowel disease (IBD) is a group of inflammatory diseases that involve chronic inflammation of all or part of the gastrointestinal track. In the US, it is estimated that about 1.5 million people currently suffer from IBD, yet the causal factors of the disease remain to be fully elucidated. In most patients, the body mounts immune responses against elements of its own digestive system. These diseases of intestine impart a significant and negative impact on the well-being of patients as well as their social environment. Understanding the underlying mechanism of the induction and perpetuation of IBD as well as the roles of immune system in IBD pathogenesis is necessary for the development of novel effective treatment and prevention strategies^1, 2^.

Experimental animal models have been very useful to address those gaps in that they can provide fundamental insights into disease-associated mechanisms of immunologic dysregulation and intestinal homeostasis^4, 5^. Several different animal models have been developed to study the pathogenesis of IBD^6, 7^. Among those, injection of an anti-CD40 agonist antibody into immune-deficient animals has been shown to induce pathogenic inflammation in the colon that depends on innate immune inflammation. This model is commonly used for studying the role of innate immune cells such as macrophages and dendritic cells in IBD pathogenesis^8^. However, using those models to study genetically susceptible loci often requires the availability of genetically modified animals for the target gene to establish a causal relationship between genetic susceptibility and IBD pathogenesis. Genome wide association analysis and meta-analysis has identified over 230 different IBD loci, and has been instructive in identifying mechanisms and pathways that are important in IBD development^3^. Identifying the functional connection between such association analysis and the key genetic driver(s) in IBD development is crucial for the discovery of effective therapeutics for patients. In this regard, it is important to note that the traditional strategy to generate knockout animals requires the generation of individual animal strain for each gene of interest, which is hardly amendable in a timely manner considering the large number of IBD-associated loci.

The CRISPR/Cas9-based genome editing system has recently emerged as a cutting-edge technology that has substantially accelerated animal model generation^9–13^ because of its potential to enable the rapid genetic manipulation of specific genes. Multiple approaches have been used to deliver CRISPR into animal models. Hydrodynamic injection, cationic lipid-mediated delivery, and adeno-associated virus (AAV)-mediated delivery successfully allow for delivery of CRISPR to adult animals with good efficiency^13–17^. However, those approaches generally only deliver CRISPR to certain regions of the body. Whole body genome editing normally involve embryos and breeding, which require specialized expertise and also take approximately a year to generate the strain with the desired genetic alteration. The modification of hematopoietic stem cells (HSCs) or Lin-Sca1+Kit+ (LSK) cells allows the fast and efficient generation of adult animals bearing desired CRISPR edits in the immune system. We hypothesized that utilizing the CRISPR/Cas9 technology with HSCs can meet the urgent need for the development of a fast and efficient system to introduce genetic edits *in vivo* for a large number of target genes.

Using CD40 as a model target, we here established a CRISPR-based methodology to ablate the expression of this key regulator in HSC-derived immune cell populations and validated our approach in the CD40 agonist-induced colitis model. We established the combined method of CRISPR/Cas9 knockout and HSC transplantation for studying a target gene in the development of IBD pathogenesis. Additionally, we demonstrated that an efficient knockout of CD40 by CRISPR/Cas9 *in vivo* significantly reduced disease induction in the experimental model. Our study demonstrated the feasibility of using this newly established methodology as a novel and robust platform to determine the functional importance of new target genes in the immune system for IBD and potentially other autoimmune diseases.

## Results

### Establishment of a transplantation protocol using Lin-Sca1+Kit+ (LSK) cells

We designed an experimental outline to establish and validate a protocol to reconstitute irradiated animals using Lin-Sca1+Kit+ (LSK) cells (Fig. 1A). First, we evaluated the methodology to enrich LSK cells by pre-treating the donor animals with 5-fluorouracil. We found that despite an increase of lin-Sca1+ cells in the donor bone marrow, c-kit expression on those cells markedly decreased (Fig. S1A). This data is consistent with prior findings showing that 5-fluorouracil treatment is associated with decreased c-kit expression^18, 19^. Therefore, we isolated LSK cells from CD45.2+ B6 animals following a two-step sorting as described previously^20^. Sorted cells were injected into lethally irradiated CD45.1+ congenic recipients. CD45.1 and CD45.2 were used as markers to distinguish donor versus recipient cells. The quality of sorted LSK cells was monitored using a mouse Colony-Forming Unit Assay *in vitro* prior to transplantation (data not shown). To monitor engraftment, five animals were taken down every two weeks after transplantation. As shown in Fig. 1B, over 80% engraftment was achieved in the spleen, bone marrow, and blood within two weeks post-transplantation. A detailed characterization of different immune cell types showed that B cells, CD11b+ macrophages and CD11c+ dendritic cells (DC) fully engrafted at week two (Fig. 1C, 1D, and Fig. S1B). In contrast, the T cell compartment required a minimum of 12 weeks to reach around 90% engraftment (Fig. 1C and 1D). This observation is in line with the fact that T cells need to undergo thymic selection to fully mature. To further ensure the proper hematopoiesis in the reconstituted animals, we examined the cell subsets within the T cell and myeloid compartment and validated the generation of regulatory T cells and neutrophils by CD25 and Ly6G staining within the T and myeloid population respectively (Fig. 1E and Fig. S1C). In summary, we established a protocol to reconstitute the murine immune system *in vivo* by transplanting LSK cells into lethally irradiated recipients.

**Fig. 1.**
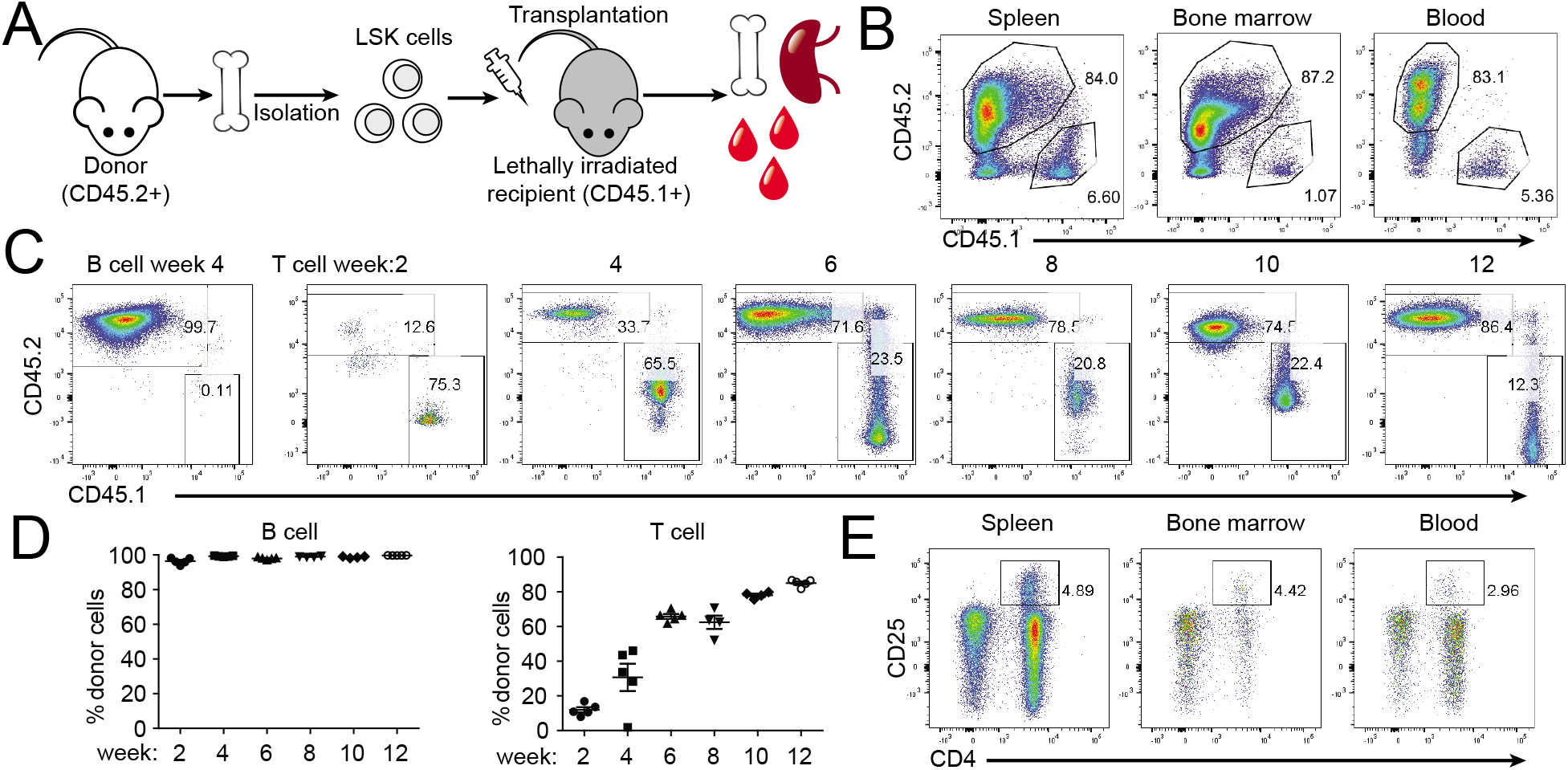
Establishment of a transplantation protocol using LSK cells. A. Experimental setup: LSK cells were isolated from CD45.2+ donor mice and then transferred to lethally irradiated CD45.1 congenic B6 animals. Every 2 weeks post-transfer, 5 mice were euthanized. Spleen, bone marrow and blood were isolated and the engraftment rate was evaluated by FACS. B, Representative FACS plots in the spleen, bone marrow and blood at week 2 post-transplantation. C. Representative FACS plots of B cells at week 4 and T cells at different timepoint post-transplantation. D. Percent of donor B and T cells at different timepoints post-transplantation. Each plot represents a data point from one animal. E. Representative regulatory T cell population in spleen, bone marrow and blood at week 12 post-transplantation. Data are showing a representative experiment (n=2, five animals per group).

### Optimization of lentiviral infection of LSK cells

Next, we optimized the lentiviral infection conditions for LSK cells. LSK cells were infected with lentiviruses encoding vexGFP, mCherry, or Thy1.1 and used to reconstitute lethally irradiated recipients. Both expression of vexGFP and mCherry, were detected 3 weeks post-transplantation while Thy1.1 was undetectable (Fig. 2A and data not shown), suggesting that vexGFP and mCherry can be used to track infected cells *in vivo*. In order to further increase the infection rate, we concentrated the virus supernatant 100-fold by ultracentrifugation and confirmed the increase in infectious units after concentration by titering the virus on 293T cells and primary human T cells. However, when comparing unconcentrated and concentrated virus preparations with LSK cells, no detectable infection was observed with concentrated virus (data not shown). Therefore, we chose to use unconcentrated virus to infect LSK cells in our following experiments.

**Fig. 2.**
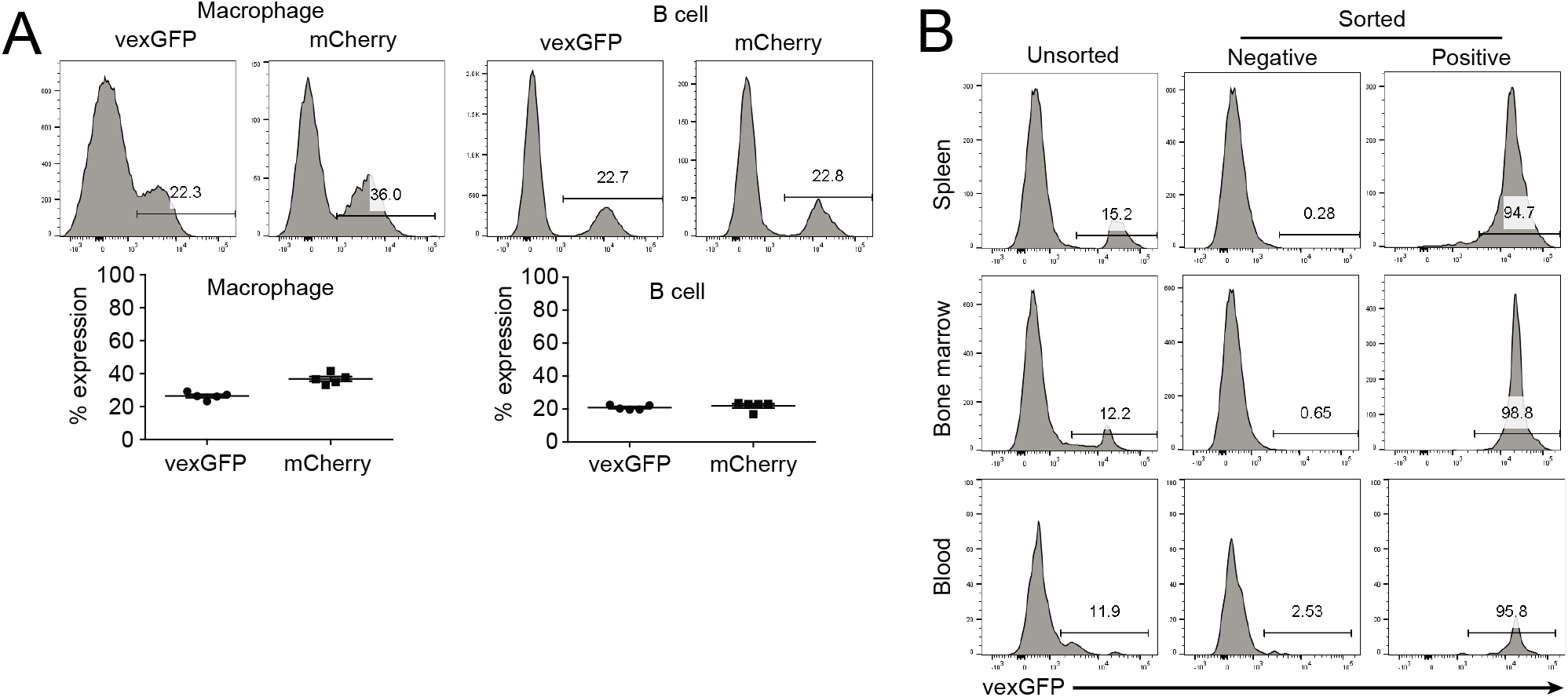
Optimization of lentiviral infection of LSK cells. A. Marker gene expression *in vivo* post-lentiviral infection. Mice are reconstituted with LSK cells infected by vexGFP or mCherry expressing lentivirus. Upper panel shows representative vexGFP and mCherry expression in macrophages and B cells. Lower panel, percent of vexGFP and mCherry from the group of treated animals. Individual dots represent data points obtained from single animals. B. Isolation of infected LSK cells. LSK cells were infected by virus encoding vexGFP at a MOI of 1. Infected cells were sorted by FACS based on vexGFP expression. Unsorted cells, negative population, and positive population from the sort were injected to recipients. Mice were harvested 3 weeks post-reconstitution. Representative FACS plots from each group are depicted. Data are showing a representative experiment (n=2, five animals per group).

In order to generate animals with more homogenous knockout of the immune system, we further purified infected cells based on marker gene expression. LSK cells were infected by vexGFP-expressing virus at a multiplicity of infection (MOI) of one, and then sorted for vexGFP after infection. In mice reconstituted using sorted donor cells, over 90% of immune cells expressed vexGFP *in vivo* (Fig. 2B). These data conclusively demonstrate that we have established a platform to infect LSK cells using lentivirus and to further purify the infected cells to use as donor cells for immune system reconstitution *in vivo*.

### CRISPR-based modulation of CD44 expression validates the novel *in vivo* platform as a tool to alter protein expression levels in hematopoietic cells

Next, we chose to use CD44, a marker gene that is expressed by many leucocyte subsets, to evaluate the editing efficiency of our platform. The sequence of guide RNA (gRNA) targeting CD44 (SgCD44) was obtained from a previously publication^21^, and we validated its efficacy *in vitro* (Fig. 3A)^21^. We generated mice using LSK cells from Cas9 knock-in mice^22^ infected with lentivirus encoding vector, scramble control (SgNone) or SgCD44, and evaluated engraftment rate in the spleen, bone marrow and blood 8 weeks post-transplantation. As shown in Fig. 3B, 3C and 3D, cells expressing SgCD44 showed a strong reduction in CD44 expression compared to cells expressing vector or SgNone, although the magnitude of reduction differed across different cell types. Thus, we have established an *in vivo* model system to edit genes of interest using CRISPR.

**Fig. 3.**
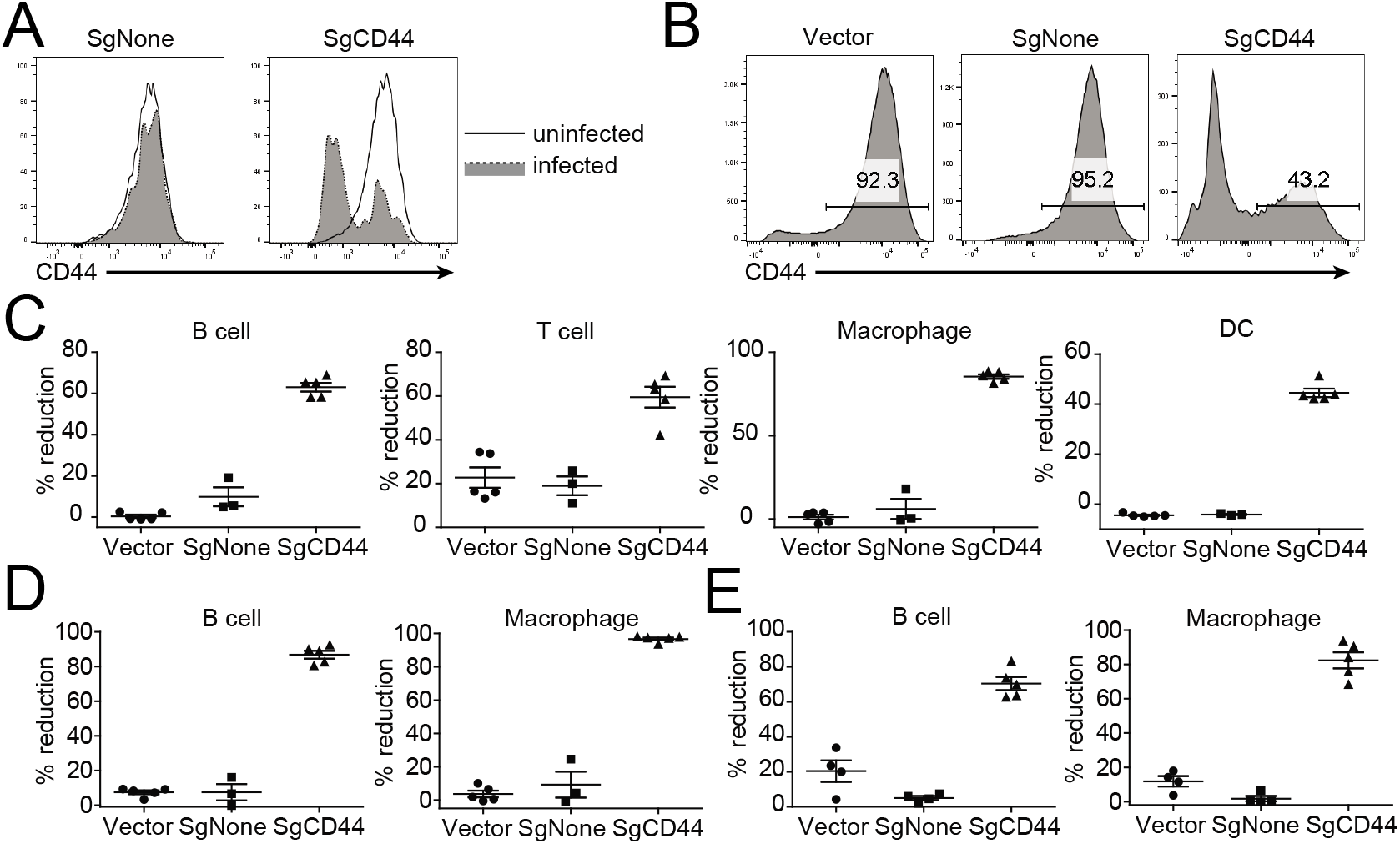
Modulation of CD44 expression using CRISPR/Cas9-based system. A. Downregulation of CD44 expression *in vitro*. Splenocytes were isolated from Cas9 KI mouse, and infected with viruses encoding a none-targeting control (SgNone) or a gRNA against CD44 (SgCD44). B, C, D. Downregulation of CD44 expression *in vivo*. Mice were generated to express the vector control, SgNone or SgCD44. Shown is representative FACS blot of CD44 expression from splenic B cells. C. Reduction of CD44 expression in B cell, T cell, macrophages and DCs in the spleen. D and E. Reduction of CD44 expression in B cells and macrophages in B cell from bone marrow (D) and blood (E). Data are showing a representative experiment (n=3, five animals per group).

### CRISPR-based knockout of CD40 ameliorates disease in a CD40 agonist-induced colitis model

We next tested combining our CRISPR-knockout strategy with an experimental model of IBD for proof of principal experiments. CD40 pathway has been associated with several types of autoimmune diseases including colitis, arthritis and lupus. CD40 agonist antibodies are known to induce systemic inflammation and colitis in mice^8, 23^. We hypothesized that knocking out CD40 using our platform will prevent disease development in the CD40 agonist-induced colitis model. To test this hypothesis, we designed three different gRNAs (SgCD40.1, SgCD40.2, and SgCD40.3) that target mouse CD40 and validated their efficiency *in vitro* using 293 cells stably expressing CD40. All three gRNAs reduced CD40 expression with an efficiency of 60 – 90% (Fig. 4A). When introduced *in vivo*, all three gRNAs achieved around 90% efficiency in impairing CD40 expression in splenic B cells (Fig. 4B). Colitis induction was performed using an anti-CD40 agonist antibody. As shown in Fig. 5, mice reconstituted with CD40 gRNA-expressing LSK cells exhibited reduced disease. All three gRNAs were able to protect the animals from weight loss and clinical manifestation measured by endoscopy score (Fig. 5B and 5C). A detailed examination of immune cell activation and infiltration revealed subtle differences in mice receiving different gRNAs: SgCD40.1 was associated with the strongest protection of disease induction; reduced CD86 upregulation and cell infiltration shown by CD3 staining for T cells. Also, IBA1 staining for myeloid cells was observed with mice receiving SgCD40.2 with lower efficiency than SgCD40.1. The impact of SgCD40.3 on myeloid and T cell infiltration was intermediate between SgCD40.1 and SgCD40.2 (Fig. 5D and 5E). The efficiency of the three gRNAs to inhibit disease induction correlated well with their efficiency to reduce CD40 expression *in vitro* in 293 cells (Fig. 4A). These data suggested that CD40 expression was efficiently ablated *in vivo* using our strategy and therefore induction of colitis was significantly reduced. In summary, we have established a novel *in vivo* CRISPR/Cas9-platform to study the functional role of target genes during colitis pathogenesis in mice.

**Fig. 4.**
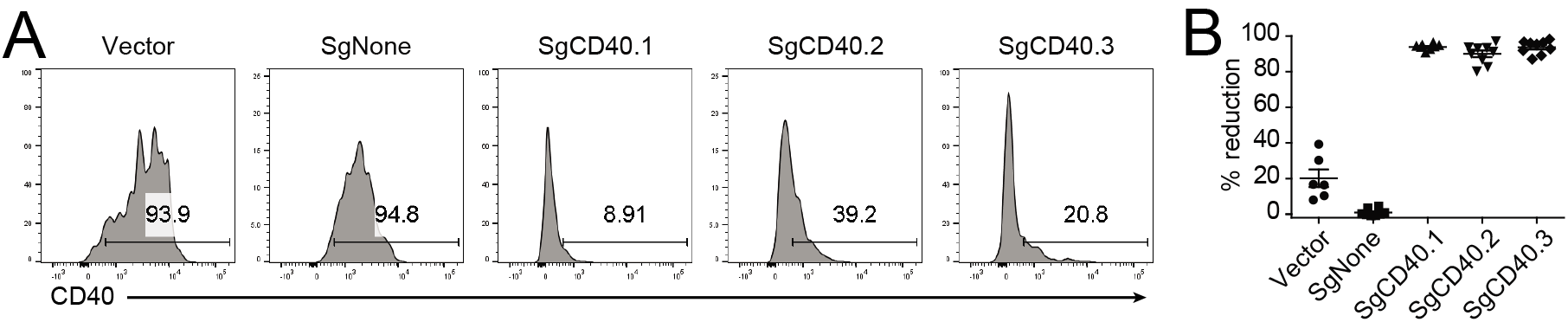
Modulation of CD40 expression using CRISPR/Cas9-based system. A. Reduction of CD40 expression by CRISPR/Cas9 *in vitro*. 293 cells stably expressing CD40 were infected with viruses encoding vector, a scramble control (SgNone), or different gRNAs for CD40. B. Reduction of CD40 expression by CRISPR/Cas9 *in vivo*. Mice were transplanted with LSK cells infected viruses encoding corresponding constructs. Expression of CD40 in spleen, bone marrow and blood was evaluated 8 weeks post-transplantation. Shown are results from splenic B cells. Data are showing a representative experiment (n=2, five animals per group).

**Fig. 5.**
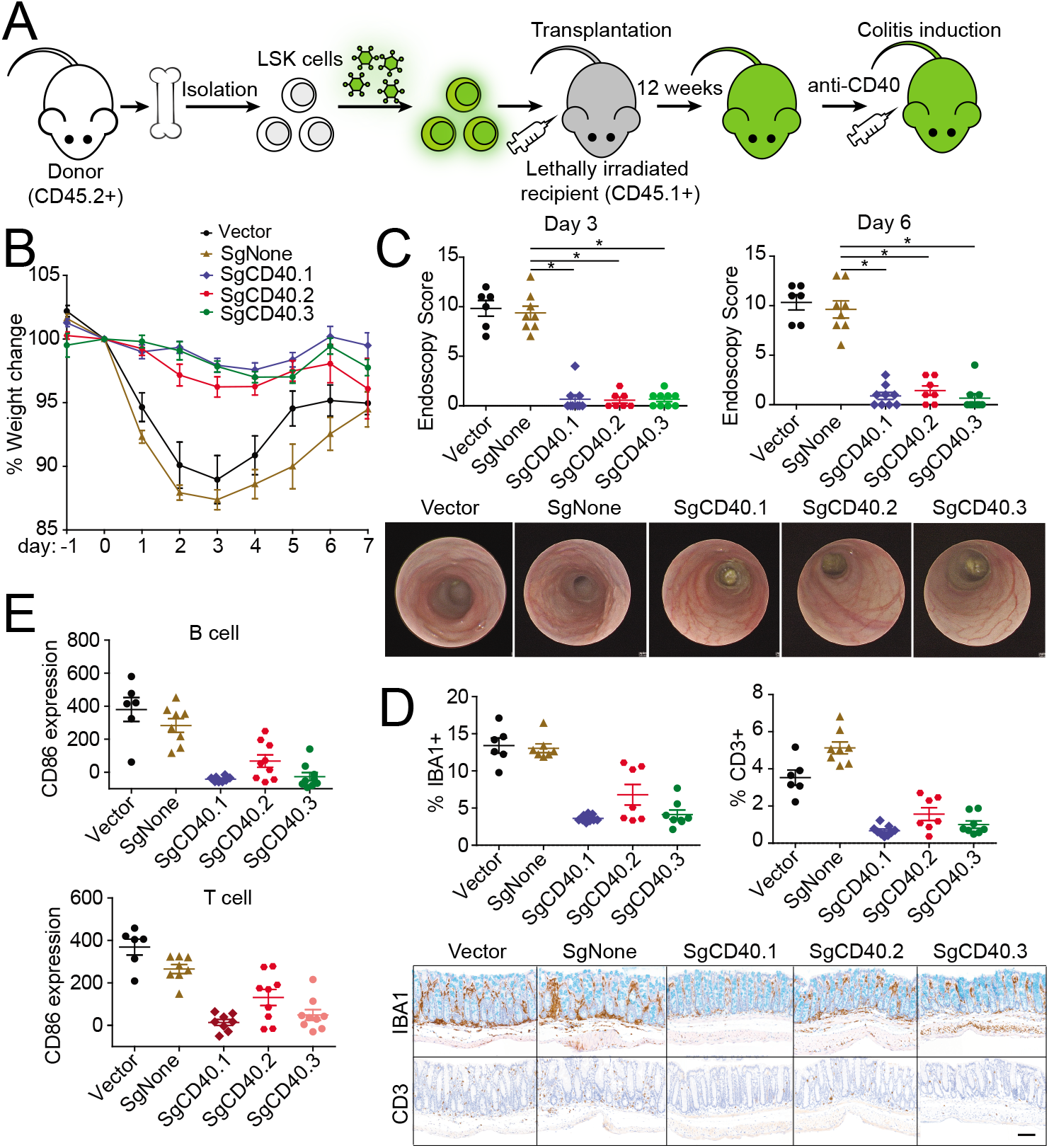
CRISPR-based knockout of CD40 ameliorates disease a CD40 agonist-induced colitis. A. Experimental procedure. LSK cells were isolated and infected with lenti-viruses expressing control or gRNAs for CD40. Infected cells were sorted out based on marker gene (vexGFP) expression and used to transplant lethally irradiated CD45.1+ recipient mice. 12 weeks post-transplantation, colitis was induced by injecting CD40 agonist antibody. Disease induction was monitored based on body weight change (B), endoscopy score at day 3 and day 6 post-anti-CD40 injection (C), percent of IBA1+ and CD3+ cell area of total mucosal area (D) and upregulation of CD86 expression in splenic B and T cells (E). Representative day 6 endoscopy and day 7 histology images are shown in C and D respectively. Scale bar, 100μM. *=*P*<0.001 by one-way ANOVA and Dunnett’s Post Hoc. Data are showing a representative experiment (n=2, ten animals per group).

## Discussion

The work presented in the current paper introduces a novel system to study gene function *in vivo* using CRISPR/Cas9-based genome editing technology. First, we optimized the protocol for generating specific knockout mice using LSK cells and, next, introduced genetic edits to LSK by lentiviral infection. By enriching infected donor cell, we achieved over 90% gene editing efficiency in the immune cells *in vivo*. As proof of concept, we demonstrated that the successful ablation of CD40 expression in the immune system inhibited colitis development in animals in which experimental IBD was induced by an anti-CD40 agonist antibody.

We first established a methodology for LSK cell isolation and transplantation in order to introduce CRISPR *in vivo*. We purified the LSK cells by two steps of cell sorting and achieved over 80% engraftment rate *in vivo* within two weeks post-transplantation. A detailed investigation at different timepoints posttransplantation revealed that, at week two, B cell and myeloid cells in the animals were mostly from donor cells, whereas it took 12 weeks for T cells to reach more than 80% engraftment. This is because T cells undergo maturation in the thymus before migrating into lymphoid organs^24^. We therefore successfully built a system to fully reconstitute the recipient animals by LSK cell transplant.

As shown previously^10, 12, 25^, CRISPR generates knockout of target gene by creating double strand DNA breaks and introducing insertion or deletions (indels) in a random fashion. The same gRNA delivered by lentiviral infection may introduce different mutations in the genomic DNA in different LSK cells; it will nevertheless induce a functional knockout in the gene if the mutation results in a shift in the open reading frame. We found that not every cell expressing vexGFP, the marker for lentiviral infection, bears knockout of the target gene. Several conditions may contribute to this scenario. Certain cells expressing the marker gene do not express gRNA due to intrinsic events during lentiviral infection; editing efficiency of gRNA is affected by Cas9 expression levels^21, 25^, chromatin accessibility and guide sequence secondary structure^26, 27^, and the editing events may not necessarily occur for each gRNA in each cell; indels do not necessarily result in frame shift in the gene and an in-frame mutation may produce a functional proteins that’s indifferent to WT protein.

It is noteworthy that the CD44 gRNA sequence in our experiments did not result in same level of editing *in vivo* in different cell subsets. A similar phenomenon has been shown previously. For example, the same gRNA demonstrated higher efficiency in reducing target gene expression in 293 cells than in pluripotent stem cells (iPSC) and mesenchymal stem cells (MSC)^28^. The used CD44 gRNA was shown to be inefficient in a previous report due to limited Cas9 expression in the system^21^. We reasoned that this CD44 gRNA was not efficient enough to achieve high levels of editing in LSK cells immediately after lentiviral infection. Since lentiviral integration results in persistent gRNA expression, the genetic editing might occur during or after the LSK cell differentiating into immune cell subsets and, therefore, result in differences depending on the cell subset. It is also plausible that CD44 is involved in the function and differentiation of immune cell subsets^29^ and ablating CD44 expression impacts cell subset maturation or migration to circulation and different lymphoid organs. Together, our data confirmed that efficiency of the gRNA varies among cell types and a screen of different gRNA sequences *in vitro* prior transplantation will be beneficial to ensure sufficient genetic editing *in vivo*.

Genetic alterations in LSKs change can only be applied to alter targets expressed in the hematopoietic lineage. We, therefore, hypothesized our system could be very useful in studying the function of genes in the immune system in the recipient animals. We therefore evaluated our platform in the anti-CD40 agonist induced colitis model, since the pathogenesis in this model is primarily driven by CD40-mediated immune cell activation. We demonstrated efficient knockout of CD40 *in vivo* using the CRISPR/Cas9 system which strongly inhibited colitis development upon CD40 agonist antibody injection. The three gRNAs targeting CD40 differed in their efficiency reducing CD40 expression in B cells. The vector control also showed slight downregulation of CD40 expression probably reflecting the extent to which modifications induced by the lentivirus affect target gene expression. Although all three gRNAs significantly ameliorated disease induction measured by body weight loss and endoscopy scores, differences were observed with regards to immune cell activation and infiltration in a manner correlated to their editing efficiency *in vivo*. Although we only demonstrated the impact of gene editing in a colitis model, it is fair to assume that this approach can be equally applied to investigate the loss of function of a target gene in other pharmacodynamic models or disease models. In addition to ablation of a gene of interest, this method may also be modified to study the gene function by overexpression or mutagenesis. Our data demonstrated a strong correlation between gRNA efficiency and their efficacy in blocking gene function, and suggested our CRISPR/Cas9-based platform can potentially be used for biology validation for novel therapeutic targets.

In summary, we here introduce a strategy that facilitates the challenging task of performing large scale of functional genomic screen *in vivo* in the context of disease models. In this regard, IBD as a diverse group of disorders with complex pathogenesis Is an ideal condition because of the plethora of regulatory genes that have been identified with genetic and genomic analysis. Our study provides a new angle to solve this issue, which is delivering a CRISPR library to pooled LSK cells to reconstitute a new immune system with most if not all immune cell subsets. We will then be able to screen for genes that provide desired cellular functional phenotype in the reconstituted animals.

Going forward, it is fair to anticipate that this work will pave the road to better characterize the importance of complex immune signaling networks in the regulation of IBD pathogenesis. The ability to validate the functional consequences of target perturbation in vivo after identification of genetic susceptibility loci or genetic association will facilitate drug development as correlative data can be substantiated with functional studies. As such, it is to be anticipated that the use of a platform as the one describe in our study will significantly accelerate the time span between basic research discoveries and clinical practice.

## Experimental procedures

### Mice

Cas9 Knock-in animals (Jackson order# 026197), C57BL/6 Ly5.1 (Jackson order# 002014), Scid (Jackson order# 001913) and C57BL/6 (Ly5.2 Jackson order #000664) mice were all ordered from Jackson Labs. All studies involving animals were performed according to protocols reviewed and approved by the Abbvie IACUC.

### Plasmids and guide RNA design

gRNA-encoding plasmids were constructed upon the lentiGuide-Puro (Addgene Plasmid #52963) vector, in which the puromycin resistant element was swapped to sequence that encodes vexGFP, mCherry or Thy1.1. The scramble control sequence or the sequence targeting CD44^21^ were subsequently cloned into the vexGFP vector. The design of the guide RNA sequences targeting CD40, and the CRISPRv2 constructs encoding the relevant gRNAs were obtained from Genscript (https://www.genscript.com/CRISPR-plasmid-gRNA-cas9.html?src=pullmenu3). The CD40 gRNA sequences are as follows: SgCD40.1, AGCGAATCTCCCTGTTCCAC; SgCD40.2, GACAAACAGTACCTCCACGA; SgCD40.3, ACGTAACACACTGCCCTAGA

### Lentivirus production

Lentiviral particles were produced as described previously^30^. Briefly, gRNA-encoding plasmid was cotransfected into 293T cells together with VSV-G and pLEX packaging plasmids. Medium was changed 18 hours after transfection, and the virus-containing supernatants were collected 24-48 h after medium exchange. For virus concentration, the harvested supernatants were centrifuged at 16500 rpm for 90 min, and the pelleted virus was resuspended in a volume that concentrated the virus 50 to 100 times.

### CRISPR knockout mice generation using Lin-Sca1+Kit+ (LSK) cells

LSK cells were isolated and infected as previously described^20^. Briefly, bone marrow cells were collected from femurs and tibias of adult animals after red blood cell lysis. CD117 (c-kit)+ cells were isolated using CD117 magnetic beads (Miltenyi) and next sorted for lin- and Sca1+ using a BD Aria sorter. In some experiments, 5-fluorouracil was injected at 150 mg/kg one week before LSK cell isolation. The pluripotency of the LSK cells isolated were confirmed using Colony-Forming Unit Assay according to the manufacturer’s instruction (STEMCELL Technologies). The LSK cells were rested overnight in SFEM medium (STEMCELL Technologies) with a mixture of cytokines (100 ng/ml Fit3L, IL-7, SCF, TPO; all from PeproTech) before transferred to Retronectin pre-coated plate (50ug/ml; Clontech). Lentivirus encoding gRNA or scramble control were added to the cells and spun down at 2000rpm for 20min. The infected cells were then cultured for 2 days before being injected into animals. In some of the experiments, the infected cells were sorted based on vexGFP expression 2 days post-infection. Recipient animals were irradiated with gamma irradiation at 600 rads twice 3 h apart. LSK cells were intravenously injected to the recipient animals 3 h after the 2^nd^ dose of irradiation. After bone marrow transplant, mice were moved to autoclaved cages and treated as immune deficient animals.

### *In vitro* efficiency evaluation of gRNAs targeting CD44 and CD40

To evaluate gRNA for CD44 *in vitro*, splenocytes from Cas9 KI mice were isolated and stimulated by 1μg/ml anti-CD3e (eBioscience) in the presence of viral particles encoding a scramble control (SgNone) or gRNA targeting CD44 (SgCD44). The cells were expanded in the presence of 100U/ml IL-2 (PeproTech) for 2 weeks before staining for surface CD44 expression.

A pcDNA3 plasmid encoding murine CD40 were transfected into 293 cells, and cell line stably expressing CD40 were selected by single cell cloning. The stable cell line were infected by lenti-viruses expressing a scramble control (SgNone) or different gRNAs targeting CD40 including SgCD40.1, SgCD40.2 and SgCD40.3. Expression of CD40 was evaluated by FACS 2 weeks post-infection.

### CD40-agonist induced colitis model

CD40 agonist antibody (FGK4.5, BioXcell) was injected intraperitoneally at 10 mg/kg per animal on day zero. Body weight was monitored every day beginning day −1. Video endoscopy on day 3 and 6 following CD40 antibody injection was performed to evaluate disease in-life. Briefly, mice were first anesthetized with 2% isofluorane, and then given an enema using DPBS to prepare the colon for endoscopy. The endoscope (Karl-Storz) was inserted into the rectum, advanced to the proximal colon, and then slowly withdrawn while taking a video recording. Images were taken at 3, 2, and 1cm distal of the anus. Each image was evaluated individually and scored to determine disease severity. On day 7, all animals were euthanized; the colon was harvested for histopathological assessment and the spleen was weighed then prepared for analysis by flow cytometry. To analyze statistical differences between control and CD40 knockout groups, One-way ANOVA’s with Dunnett’s Post Hoc test (vs SgNone control group) were performed to determine minimum significance of p<0.05 using GraphPad Prism 5.0 (GraphPad Software, San Diego, CA).

### Immunohistochemistry (IHC)

Formalin-fixed and paraffin wax-embedded (FFPE) tissue sections of mouse colon were used for immunohistochemical staining of ionized calcium binding adaptor molecule 1 (IBA1) and CD3. Immunohistochemical staining was performed on a Leica Bond RX^®^ automated Stainer (Leica Biosystems). The Bond Polymer Refine DAB Detection kit (Leica Biosystems, DS9800 DAB) was used and all procedures followed manufacture’s protocols. The sections were subjected to ethylenediaminetetraacetic acid (EDTA) based antigen retrieval for 20 min followed by application of the primary antibody. For identification of macrophages, antibody to IBA1 (Wako Chemicals, USA) was used at 0.15μg/ml for 30 min at room temperature. To identify T-lymphocytes, anti-CD3 antibody (Lab Vision, Fremont, CA) was used at 0.08μg/ml for 15 min at room temperature. Following staining samples were counterstained and cover slipped for image analysis.

### FACS

Tissues including spleen, bone marrow and blood were isolated from animals, and cells were extracted from the tissues. Red blood cells were lysed by lysis buffer (eBioscience) and the cells were stained using fixable live/dead dye (Invitrogen) according manufacture’s protocol before stained with antibodies of interest at 4°C for 20min. Cells were washed after staining and analyzed by a BD LSRFortessa™ cell analyzer. Antibodies used include: B220, GR-1, Ter119, CD11b, CD3, CD40, CD44, CD45.1, CD45.2, Thy1.1, and Sca-1 (all from Biologend).

---

**Supplementary Fig. 1.**
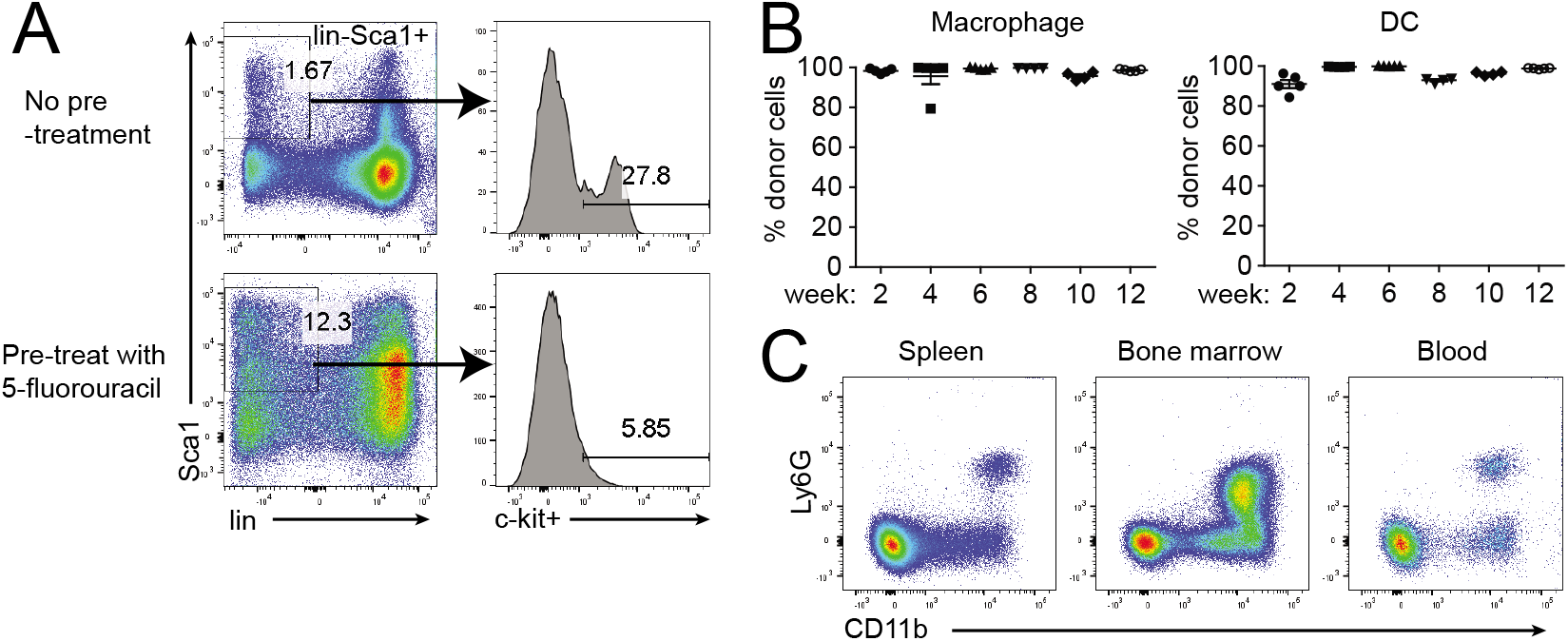
Optimization of LSK cell transplantation protocol. A. Impact of 5-fluorouracil in LSK cell enrichment. Mice were treated or not treated with 5-fluorouracil one week before LSK cell isolation. B. Neutrophil in the spleen, bone marrow and blood from LSK cell reconstituted mice at week 12 post-transplantation. Representative image from the corresponding group of mice is shown. C. Percent of donor cells in macrophage and DCs in LSK cell reconstituted mice at different timepoints after transplantation. Data are showing a representative experiment (n=2, five animals per group).

## Author contributions

Rui Wang designed majority of the experiments, interpretation of the results, and execution of some of the experiments. Ning Sun contributed to the plasmid design and generation. Sean Graham and Ruoqi Peng contributed to the execution of the in vivo studies. Xiaochun Zhu contributed to the in vitro studies including lentiviral infection of cells. Jamie Erikson and Robert Dunstan contributed to the IHC analysis. Marc Wurbel, Namjin Chung, Edda Friebiger, Tariq Ghayur and Jijie Gu contributed to the evaluation of the work and preparation of the manuscript.

